# MOHCA-seq: RNA 3D models from single multiplexed proximity-mapping experiments

**DOI:** 10.1101/004556

**Authors:** Clarence Yu Cheng, Fang-Chieh Chou, Wipapat Kladwang, Siqi Tian, Pablo Cordero, Rhiju Das

**Affiliations:** Department of Biochemistry, Stanford University, Stanford, California 94305, United States; Biomedical Informatics Program, Stanford University, Stanford, California 94305, United States; Department of Physics, Stanford University, Stanford, California 94305, United States

## Abstract

Large RNAs control myriad biological processes but challenge tertiary structure determination. We report that integrating <u>m</u>ultiplexed •<u>OH</u> <u>c</u>leavage <u>a</u>nalysis with tabletop deep <u>seq</u>uencing (MOHCA-seq) gives nucleotide-resolution proximity maps of RNA structure from single straightforward experiments. After achieving 1-nm resolution models for RNAs of known structure, MOHCA-seq reveals previously unattainable 3D information for ligand-induced conformational changes in a double glycine riboswitch and the sixth community-wide RNA puzzle, an adenosylcobalamin riboswitch.

## Main Text

RNAs form and interconvert between specific secondary and tertiary structures to perform a wealth of essential biological functions^1,2^. Conventional high-resolution structure determination techniques provide critical insights into RNA behavior but are limited in throughput and are challenged by the large sizes and conformational heterogeneity of many RNAs^3^. Chemical mapping techniques, such as hydroxyl radical footprinting and SHAPE, comprise an important class of fast experiments for probing RNA structure^3^ which can guide RNA structure inference with computational modeling algorithms. Nevertheless, though recent advances have enhanced the precision of chemical mapping data analysis^4–9^, these techniques do not generally give sufficient information to model folds in three dimensions.

RNA structure modeling would benefit from pairwise information for nucleotides that are proximal in the 3D fold but not necessarily Watson-Crick base-paired. Previously, we reported a technique for discovering such RNA tertiary contacts, termed Multiplexed hydroxyl radical (•OH) Cleavage Analysis (MOHCA)^10^. MOHCA relies on localized generation of hydroxyl radicals from sources that are randomly incorporated into the RNA backbone. These radicals cause strand scission events that are spatially correlated with the location of the radical source. Determining the sequence positions of the 5′- and 3′-ends of the resulting RNA fragments yields rich pairwise proximity information, which can be used to constrain computational modeling of the RNA. The original MOHCA protocol (‘MOHCA-gel’) achieved 13 Å resolution, well under the 23 Å diameter of an RNA helix, when used to model the P4-P6 domain of the *Tetrahymena* ribozyme^10^. However, MOHCA-gel required specially synthesized 2′-NH_2_-2′-deoxy-α-thio-nucleotide triphosphates for radical source attachment and on-demand backbone scission; a customized two-dimensional gel electrophoresis setup; and radioactive labeling for the readout^10^. These factors prevented MOHCA-gel from entering routine use, despite its unique ability to resolve 3D folds for any RNA.

We hypothesized that the recent availability of tabletop deep sequencing would enable a simplified, accelerated, and higher-precision MOHCA protocol, which we term MOHCA-seq. Figure 1a shows the new workflow. After randomly incorporating 2′-NH_2_-2′-dATP during transcription and coupling to isothiocyanobenzyl-Fe(III)•EDTA (both commercially available), RNAs are folded and then fragmented by activating the Fenton reaction at the backbone-tethered Fe(III) atoms using ascorbate as a reducing agent^10^. A pre-adenylated DNA adapter is ligated to the 3′-ends of the resulting fragments, permitting reverse transcription with primers harboring an Illumina adapter and experiment-specific barcodes^11^. Reverse transcription is terminated by the radical source or other correlated oxidative damage sites, so that the 5′- and 3′-ends of the resulting cDNA fragment report two positions that were proximal to (or coincide with) the radical source. A second Illumina adapter is then ligated to the cDNA, permitting paired-end sequencing on a standard Illumina platform^6,7^. MAPseeker software analysis then quantifies the data and generates a pairwise proximity map (Methods and Supplementary Fig. 1). Unlike MOHCA-gel, MOHCA-seq does not require specially synthesized nucleotides or 2D gel electrophoresis and enables detection of previously-invisible double oxidative damage events. The protocol’s throughput is further enhanced by its highly parallel nature, which we leveraged to coload multiple RNAs on single sequencing runs (Methods).

**Figure 1.**
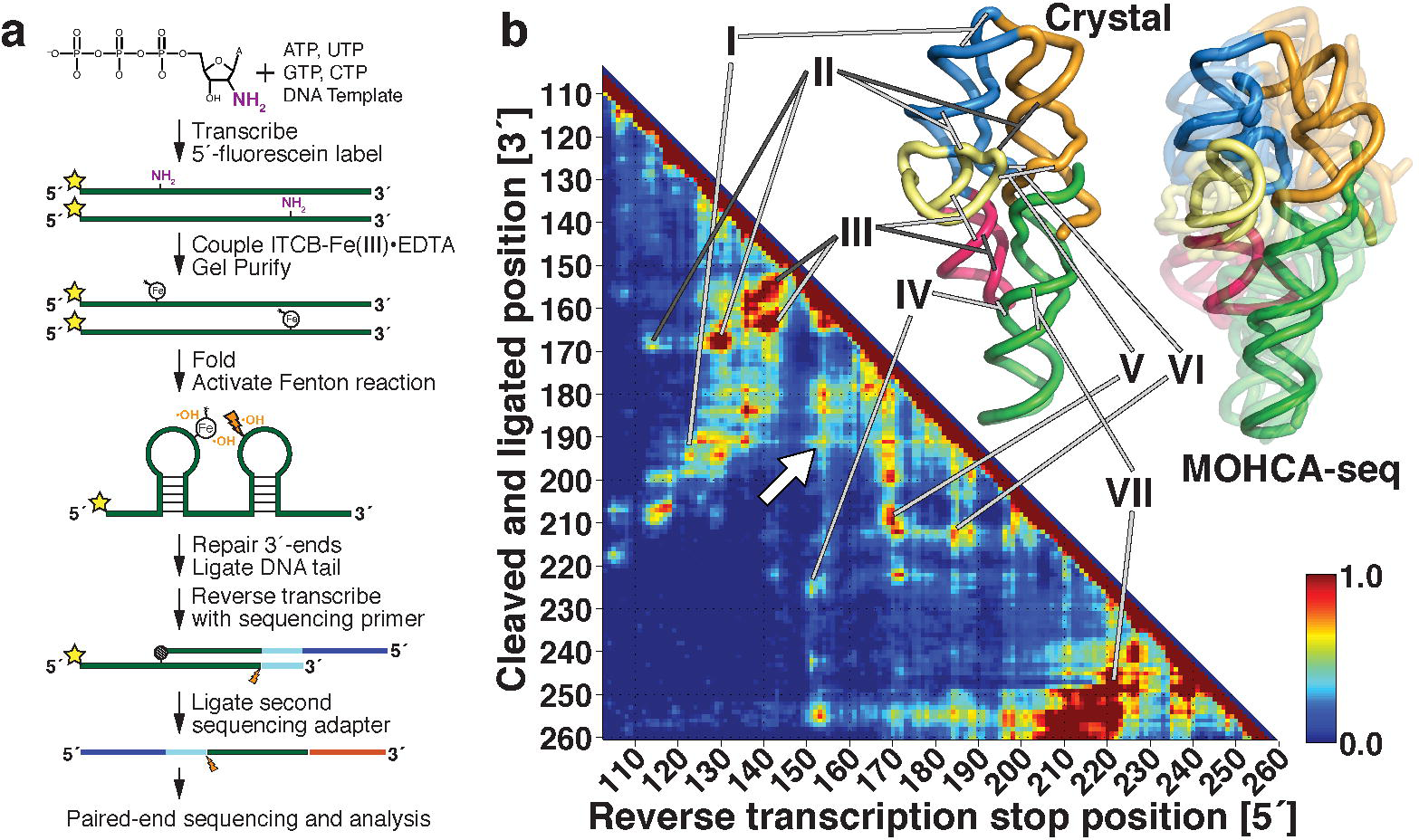
MOHCA-seq workflow and experimental validation on *Tetrahymena* P4-P6 domain. **(a)** MOHCA-seq workflow (see Methods). **(b)** The analyzed sequencing data are represented on a proximity map based on the frequency of co-occurrence of the 5′ (x-axis) or 3′ (y-axis) ends of the RNA fragments. Evidence for proximal regions of P4-P6 are provided by MOHCA-seq: (I) at the J5/5a hinge; (II) between P5c and P5a (light gray) and P5c and P4 (dark gray); (III) across the major (dark gray) and minor (light gray) grooves of P5b, (IV) between the L5b tetraloop and J6a/6b tetraloop receptor, (V) between L5c and P4, (VI) between the A-rich bulge and P4, and (VII) along the P6 helix. Some hits (white arrow) may be due to alternative conformations or radical source linker flexibility and are not represented in the crystal structure (PDB ID 1GID). Overlaid models of P4-P6 (right) are produced by *de novo* modeling in Rosetta, incorporating MOHCA-seq constraints along with solution hydroxyl radical footprinting and secondary structure inferred from prior mutate-and-map analysis. Models shown include cluster center (opaque, accuracy of 8.6 Å RMSD from crystal structure) and four other models from the largest cluster (translucent to show precision, 11.7 Å intra-cluster RMSD).

We first validated the MOHCA-seq protocol by applying it to a well-studied model system with known crystallographic structure, the 158-nucleotide P4-P6 domain of the *Tetrahymena* ribozyme^12^ (Fig. 1b, Supplementary Table 1). Hits are visible at all known secondary structure elements, with features for each helix corresponding to radicals diffusing across the major and minor grooves (Fig. 1b). Furthermore, tertiary contacts, including the bend at the J5/5a hinge, the A-minor contact between the A-rich bulge and P4, and the docking of the L5b tetraloop into the J6a/6b receptor, appear as punctate features (Fig. 1b, Supplementary Fig. 2a). Similar features were observed when we varied the radical source incorporation rate, the nucleotide at which the radical sources were attached (A, C, G, or U), and the fragmentation reaction time (Supplementary Figs. 3–5). Weak signals in the MOHCA-seq proximity maps (white arrow, Fig. 1b) that are not predicted by the crystal structure may be caused by solution flexibility of the RNA or of the ∼11 Å benzyl-thiourea linker tethering the radical source to the RNA backbone. Unlike prior MOHCA-gel work^10^, the signal-to-noise afforded by high-throughput sequencing enabled a single A-modified experiment to report on all crucial contacts in the RNA.

We expected these MOHCA-seq data to enable 3D computational modeling of the P4-P6 domain without crystallographic information. As in prior work, we used all data, including weak features not explained by the crystal structure (Supplementary Table 3), anticipating that the modeling would automatically determine the maximal subset of MOHCA-seq contacts consistent with each other and with a single fold. Indeed, application of MOHCA-seq data as pseudo-energy constraints in the *de novo* FARFAR algorithm and clustering gave the model ensemble in Figure 1b (Methods, pseudo-energy potential in Supplementary Fig. 6). The modeling achieved a precision of 11.7 Å (automatically estimated as the all-heavy-atom RMSD required to cluster 1/6 of the lowest energy models^13^) and an accuracy of 12.3 Å (median RMSD of top cluster members to crystal structure). The resolution was slightly better than the 13 Å accuracy of MOHCA-gel, which was achieved at substantially greater experimental expense^10^ (Supplementary Table 3).

After initial validation on the P4-P6 domain, we performed MOHCA-seq on three additional benchmark cases with known structures but distinct functions: the 161-nucleotide ligand-binding domain of the *F. nucleatum* double glycine riboswitch bound to two glycines^14^, the 168-nucleotide *S. thermophilum* adenosylcobalamin (AdoCbl) riboswitch bound to AdoCbl^15^, and the 120-nucleotide *in-vitro*-evolved class I ligase^16^ (Fig. 2a–c, Supplementary Fig. 2b–d, and Supplementary Table 1). In all cases, secondary structures were first defined at nucleotide resolution through mutate-and-map (M^2^) analysis (prior work^4^ and Supplementary Figs. 7 and 9). We achieved a precision of 7.1 Å and an accuracy of 10.3 Å to the crystallographic structure^14^ for the glycine riboswitch and a precision of 9.8 Å and an accuracy of 12.4 Å to the crystallographic structure^15^ for the AdoCbl riboswitch (Supplementary Table 3). Our previous modeling for the AdoCbl riboswitch during the sixth community-wide RNA-puzzle challenge was unable to disambiguate between a range of 3D folds (http://paradise-ibmc.u-strasbg.fr/rnapuzzles/6_index.html). Here, MOHCA-seq-guided modeling converged unambiguously (Supplementary Fig. 10). For the class I ligase, we achieved a precision of 8.2 Å and an accuracy of 15.4 Å to the crystallographic structure (Fig. 2c; clusters with knots were not accepted, see Supplementary Fig. 8 and Supplementary Table 3)^16^. Overall, our studies on four RNAs of known structure confirmed the ability of MOHCA-seq to resolve folds at nanometer resolution (precisions of 7–12 Å and accuracies of 10–15 Å), including cases that challenged prior approaches.

**Figure 2.**
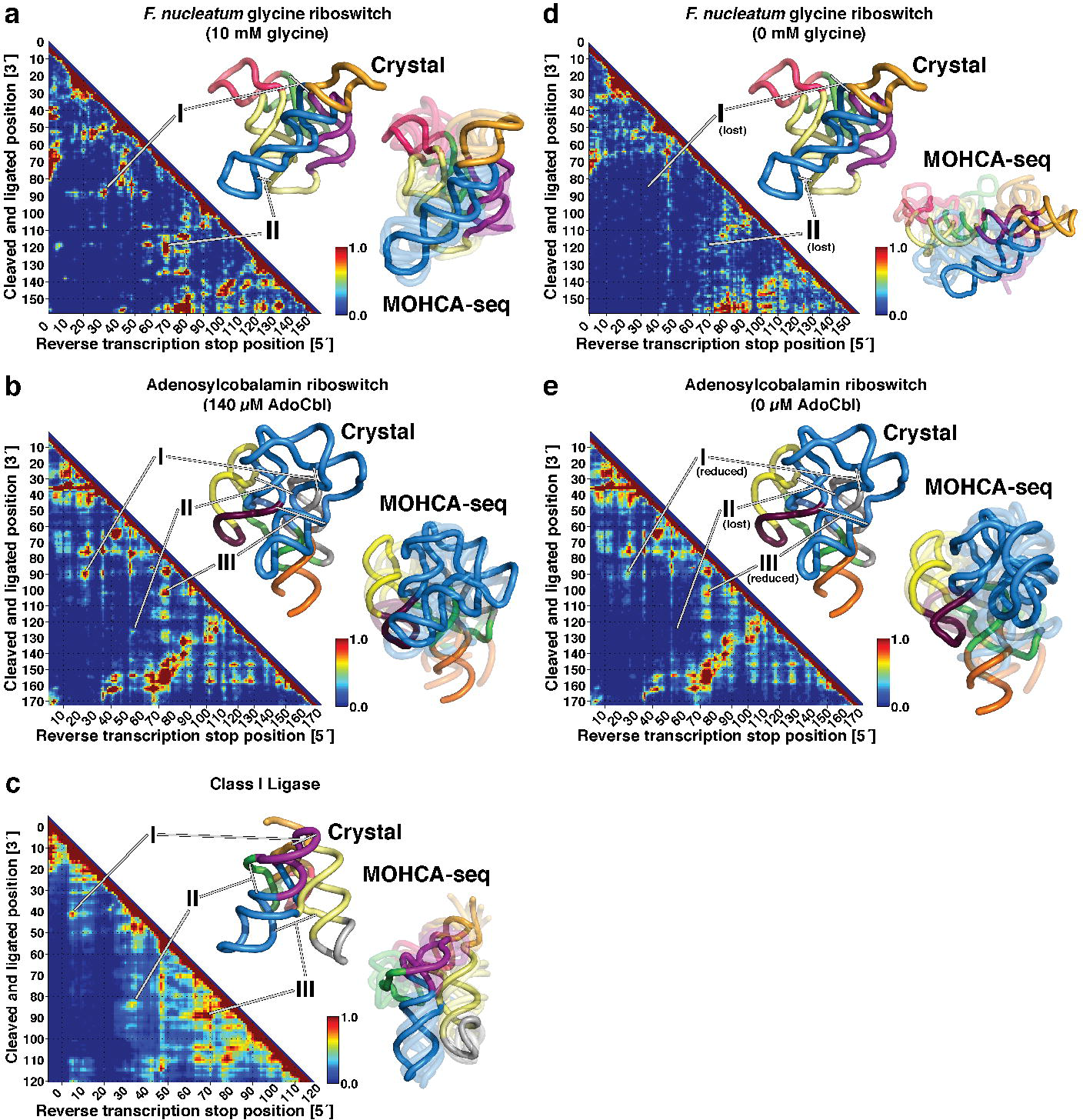
MOHCA-seq allows rapid tertiary structure mapping and modeling of varied RNAs. **(a-e)** MOHCA-seq proximity maps with crystal structures and models from Rosetta modeling with MOHCA-seq constraints shown next to each map. Models shown are the cluster center (opaque) and other models (translucent) from the largest cluster (a, c, d) or cluster centers from three modeling setups (b, e) (see Methods). **(a)** *F. nucleatum* double glycine riboswitch binding domain (PDB ID 3P49) with 10 mM glycine showing proximity of (I) P3 of aptamer 1 to J1/2 of aptamer 2 and (II) P1 of aptamer 1 to J3a/3b of aptamer 2. **(b)** *S. thermophilum* adenosylcobalamin (AdoCbl) riboswitch (PDB ID 4GXY) with 140 *µ*M AdoCbl, showing proximity of (I) P8 to P2, (II) J11-10 to L4, and (III) J8-10 to P6. **(c)** Class I ligase (PDB ID 3HHN) showing proximity of (I) P2 to J3/4, (II) J1/3 to P6, and (III) P5 to P7. **(d)** Glycine riboswitch with 0 mM glycine, showing loss of inter-domain contacts (I and II). At right, models of the unbound riboswitch adopt extended conformations, though the two aptamers fold separately. **(e)** AdoCbl riboswitch with 0 *µ*M AdoCbl, showing weakening but not complete loss of tertiary contacts, particularly between peripheral (P6-P11) and core (P1-P5) regions.

We next sought to demonstrate the ability of MOHCA-seq to infer unsolved 3D structures by probing two test cases for which crystallographic data were not available: the unliganded states of the glycine and adenosylcobalamin riboswitches. First, we probed the double glycine riboswitch in the absence of glycine. Previously proposed models of cooperativity^14^ suggested that glycine binding would be coupled to tertiary assembly of the RNA’s two domains, but previous M^2^ data could only resolve secondary structure and showed no changes upon glycine binding^4^. Strikingly, MOHCA-seq data for the ligand-free riboswitch confirmed the predicted loss of inter-domain tertiary contacts (Fig. 2d; difference map in Supplementary Fig. 11) compared to the ligand-bound riboswitch (Fig. 2a). The modeling gave a heterogenous structural ensemble with no contacts between the two glycine-binding domains, supporting the prior model (Fig. 2d).

As a second test case, we probed the ligand-free state of the AdoCbl riboswitch. Surprisingly, we observed reduction but not complete loss of MOHCA-seq hits corresponding to tertiary structure around the ligand-binding site (Fig. 2e; difference map in Supplementary Fig. 12). Modeling with the ligand-free MOHCA-seq data produced a structural ensemble whose members gave RMSDs to the ligand-bound crystal structure of 13.7 Å to 17.3 Å (Fig. 2e, Supplementary Fig. 10c, and Supplementary Table 3). These data were consistent with the formation of a conformational ensemble that samples the ligand-bound state within the sub-helical resolution of the MOHCA-seq method. Taken with available data on these and other riboswitches (ref. 17 and refs. therein), the MOHCA-seq results demonstrate that riboswitches show different levels of partial structure in the absence of ligand and must be assessed on a case-by-case basis. By enabling modeling of unliganded states, MOHCA-seq provides opportunities for detailed structure-function analyses comparing these models to thermodynamic folding measurements and probing the interplay of ligand-binding segments and flanking sequences^18^.

The MOHCA-seq method provides a rapid, automated route for modeling 3D structures of RNAs and for thereby testing and generating novel biological hypotheses. An emerging generation of high-throughput chemical mapping techniques has focused on base-pair-level RNA secondary structure, and MOHCA-seq advances this armamentarium of methods by delineating the pairwise through-space proximities needed for 3D modeling. Quantitative relationships correlating MOHCA signal intensity to structural features should further improve modeling accuracy; growth in sequencing and computing power should expand MOHCA-seq applications to larger transcripts; and development of cell-permeable MOHCA-seq nucleotide analogs may enable *in vivo* 3D structure determination. We hope that the public availability of our current and future MOHCA-seq data will help accelerate these important developments.

## Methods and Supplementary Information

A full description of the protocol for MOHCA-seq analysis, data set deposition IDs, Supplementary Figures 1–12, and Supplementary Tables 1–3 are available in the supplementary information. References 19–26 appear in the supplementary information.

## Acknowledgements

We thank members of the Das lab, V. Risca, and W. Greenleaf for discussions. We acknowledge financial support from the National Institutes of Health (5 T32 GM007276 to C.Y.C.; R01 GM102519 to R.D.), the Burroughs-Wellcome Foundation (CASI to R.D.), Bio-X and HHMI international fellowships (F.C.C.), a Stanford Graduate Fellowship (S.T.), and a Conacyt fellowship (P.C.).

### Author contributions

C.Y.C. and R.D. conceived the experiments. C.Y.C. and W.K. developed and performed MOHCA-seq experiments. S.T. performed mutate-and-map experiments. R.D. developed COHCOA analysis. P.C. developed LAHTTE analysis. C.Y.C., W.K., and R.D. analyzed the data. F.C.C. performed Rosetta modeling. R.D., C.Y.C., F.C.C., and P.C. wrote the paper. All authors discussed the results and commented on the manuscript.

## References

1. Atkins, J.F., Gesteland, R.F. & Cech, T. (Cold Spring Harbor Laboratory Press, Cold Spring Harbor, N.Y.; 2011).

2. Breaker, R.R. Cold Spring Harbor Perspectives in Biology 4 (2012).

3. Weeks, K.M. Current Opinion in Structural Biology 20, 295–304 (2010).

4. Kladwang, W., VanLang, C.C., Cordero, P. & Das, R. Nature Chemistry 3, 954–962 (2011).

5. Ding, F., Lavender, C.A., Weeks, K.M. & Dokholyan, N.V. Nature Methods 9, 603–608 (2012).

6. Mortimer, S.A., Trapnell, C., Aviran, S., Pachter, L. & Lucks, J.B. Current Protocols in Chemical Biology 4, 275–297 (2012).

7. Seetin, M.G., Kladwang, W., Bida, J.P. & Das, R. Methods in Molecular Biology 1086, 95–117 (2014).

8. Yoon, S. et al. Bioinformatics 27, 1798–1805 (2011).

9. Karabiber, F., McGinnis, J.L., Favorov, O.V. & Weeks, K.M. RNA 19, 63–73 (2013).

10. Das, R. et al. Proceedings of the National Academy of Sciences of the United States of America 105, 4144–4149 (2008).

11. Ingolia, N.T., Brar, G.A., Rouskin, S., McGeachy, A.M. & Weissman, J.S. Nature Protocols 7, 1534–1550 (2012).

12. Cate, J.H. et al. Science 273, 1678–1685 (1996).

13. Das, R., Karanicolas, J. & Baker, D. Nature Methods 7, 291–294 (2010).

14. Butler, E.B., Xiong, Y., Wang, J. & Strobel, S.A. Chemistry & Biology 18, 293–298 (2011).

15. Peselis, A. & Serganov, A. Nature Structural & Molecular Biology 19, 1182–1184 (2012).

16. Bergman, N.H., Lau, N.C., Lehnert, V., Westhof, E. & Bartel, D.P. RNA 10, 176–184 (2004).

17. Baird, N.J. & Ferre-D’Amare, A.R. RNA 19, 167–176 (2013).

18. Sherman, E.M., Esquiaqui, J., Elsayed, G. & Ye, J.D. RNA 18, 496–507 (2012).

19. Balasubramanian, B., Pogozelski, W.K. & Tullius, T.D. Proceedings of the National Academy of Sciences of the United States of America 95, 9738–9743 (1998).

20. Pogozelski, W.K. & Tullius, T.D. Chemical Reviews 98, 1089–1108 (1998).

21. Cameron, V. & Uhlenbeck, O.C. Biochemistry 16, 5120–5126 (1977).

22. Wang, B.-C. Methods Enzymol. 115, 90–112 (1985).

23. Kladwang, W., VanLang, C.C., Cordero, P. & Das, R. Biochemistry 50, 8049–8056 (2011).

24. Aviran, S. et al. Proceedings of the National Academy of Sciences of the United States of America 108, 11069–11074 (2011).

25. Cordero, P., Kladwang, W., VanLang, C.C. & Das, R. Methods in Molecular Biology 1086, 53–77 (2014).

26. Hajdin, C.E. et al. Proceedings of the National Academy of Sciences of the United States of America 110, 5498–5503 (2013).

